# Quantitative proteomics of cerebrospinal fluid from African Americans and Caucasians reveals shared and divergent changes in Alzheimer’s disease

**DOI:** 10.1101/2022.12.07.519393

**Authors:** Erica Modeste, Lingyan Ping, Caroline M. Watson, Duc M. Duong, Eric B. Dammer, Erik C.B. Johnson, Blaine R. Roberts, James J. Lah, Allan I. Levey, Nicholas T. Seyfried

## Abstract

Despite being twice as likely to get Alzheimer’s disease (AD), African Americans have been grossly underrepresented in AD research. While emerging evidence indicates that African Americans with AD have lower cerebrospinal fluid (CSF) levels of Tau compared to Caucasians, other differences in AD CSF biomarkers have not been fully elucidated. Here, we performed unbiased proteomic profiling of CSF from African Americans and Caucasians with and without AD to identify both common and divergent AD CSF biomarkers. Multiplex tandem mass tag-based mass spectrometry (TMT-MS) quantified 1,840 proteins from 105 control and 98 AD patients of which 100 identified as Caucasian while 103 identified as African American. Consistent with previous findings, the increase of Tau levels in AD was greater in Caucasians than in African Americans by both immunoassay and TMT-MS measurements. Network analysis organized the CSF proteome into 14 modules associated with brain cell-types and biological pathways. CSF modules which included 14-3-3 proteins (YWHAZ and YWHAG), demonstrated equivalent disease-related elevations in both African Americans and Caucasians with AD, whereas other modules demonstrated more profound disease changes within race. Modules enriched with proteins involved with glycolysis and neuronal/cytoskeletal proteins, including Tau, were more increased in Caucasians than in African Americans with AD. In contrast, a module enriched with synaptic proteins including VGF, SCG2, and NPTX2 was significantly lower in African Americans than Caucasians with AD. Using a targeted proteomic approach (selected reaction monitoring) followed by a receiver operating characteristic curve (ROC) analysis we measured levels of VGF, SCG2, and NPTX2, which were significantly better at classifying African Americans than Caucasians with AD. Collectively, our findings provide insight into additional protein biomarkers and pathways reflecting underlying brain pathology that are shared or differ by race.

## INTRODUCTION

African Americans are almost twice as likely to have Alzheimer’s disease (AD) compared to Caucasians (1-3). Current evidence suggests that this difference in risk could be explained by a multitude of factors including genetic ancestry and disparities in health, socioeconomic and environmental conditions (4-7). For example, genome wide association studies (GWAS) show that *ABCA7* gene has stronger associations with AD risk in individuals with African ancestry than in individuals with European ancestry (6,7). *ABCA7* also has a stronger effect size in African Americans than even the strongest genetic risk factor gene for AD, the *APOE* epsilon 4 allele (*APOE* ε4) (6). Yet, despite the *APOE* ε4 allele being more prevalent amongst African Americans, *APOE* ε4 confers a lower risk for AD compared to Caucasians (8). Beyond genetic ancestry, chronic health conditions associated with higher risk for dementia, such as cardiovascular disease and diabetes, also disproportionally affect African Americans (4,5). Furthermore, societal and environmental disparities that disproportionately affect African Americans, including lower levels and quality of education, higher rates of poverty, and greater exposure to adversity and discrimination, increase risk for both chronic diseases and dementia (4,5). This highlights how racial differences in AD risk cannot be exclusively explained by genetics alone (4). Currently, there is a gap in knowledge of the racial differences underlying pathophysiology related to AD. Therefore, a better understanding of these mechanisms can help move towards a more precise definition of AD across diverse racial, ethnic, and genetic backgrounds.

Amyloid-beta_1-42_ (Aβ_42_) and Tau, two core cerebrospinal fluid (CSF) protein biomarkers comprise the major pathologic hallmarks of senile plaques and neurofibrillary tangles found in AD brain (9,10). Thresholds between CSF Aβ_42_ and total Tau levels (tTau), established from predominantly Caucasian populations, are widely used as diagnostic measures for AD in the clinic (11-15). Emerging evidence suggests that race is associated with the levels of AD biomarkers in CSF (16,17). African Americans with AD have lower levels of CSF tau compared to Caucasians (16,17), which may explain why African Americans are diagnosed at a later disease stage (18). Consistently, African Americans have reduced rates of enrollment in clinical trials which utilize tTau/Aβ_42_ CSF ratio as an enrollment criterion (19). It is thought that elevations in CSF Tau are associated with increasing neuronal damage and cognitive impairment (20), which suggests that African Americans are symptomatic with AD even with lower levels of neuronal loss. This supports a hypothesis that there are other physiological differences contributing to the increased susceptibility and cognitive impairment in African Americans that is reflected in the CSF. An unbiased analysis into the CSF proteome of African Americans could provide insight into additional biomarkers reflecting underlying brain pathology that differ by race in AD.

Proteins are the ideal markers for understanding diseases such as AD because they are most proximal to the phenotypic changes. Unbiased proteomics of human brain coupled with network analysis has emerged as a valuable tool for organizing complex unbiased proteomic data into groups or “modules” of co-expressed proteins that reflect various biological functions linked to AD (21-24). The direct proximity of CSF to the brain presents a strong rationale to integrate the brain and CSF proteomes to identify biofluid biomarkers associated with brain pathophysiology of AD. Indeed, we recently performed an integrated human AD brain and CSF proteome analysis to reveal that approximately 70% of the CSF proteome overlapped with the brain proteome (25). While AD brain proteomic networks from large cohorts have been examined (23,26), AD CSF proteomic networks from large cohorts that include racially diverse subjects have been largely unexplored.

In this study, we used a tandem mass tag mass spectrometry (TMT-MS) approach to generate a deep CSF proteome, without albumin depletion, from over 200 control and AD samples, with over half of the samples derived from African Americans. Over 1,800 CSF proteins were organized into 14 modules based on co-expression network analysis. These modules were associated with brain cell-types and biological pathways known to be altered in AD brain, including synaptic, immune and metabolic processes. Notably, Tau mapped to a module with a magnitude of increase greater in Caucasians than African Americans with AD. In addition, network analysis revealed a core class of CSF markers that demonstrated equivalent disease-related elevations in both African Americans and Caucasians with AD, whereas other modules demonstrated more profound disease changes within race. Namely, a module enriched with post-synaptic neuronal proteins was significantly lower in level in African Americans than Caucasians with AD. Using a targeted MS approach, selected reaction monitoring (SRM), we measured the neuronal proteins within this module including VGF, SCG2, and NPTX2 which were better classifiers for AD in African Americans than Caucasians. Overall, these results demonstrate the utility of a systems-based approach in the identification of CSF proteins that could serve as markers for AD across a more diverse population. Moreover, these data highlight a need for further investigations into how AD heterogeneity varies across different racial backgrounds.

## MATERIALS AND METHODS

### CSF samples

All CSF samples were collected as part of ongoing studies at Emory’s Goizueta Alzheimer’s Disease Research Center (ADRC) including participants in the ADRC Clinical Core, the Emory Healthy Brain Study, and the ADRC-affiliated Emory Cognitive Neurology Clinic. All participants provided informed consent under protocols approved by Emory University’s Institutional Review Board. Clinical diagnosis of AD as well as classification as cognitively normal controls was based on review of clinical history, neurological examination, detailed cognitive testing, and diagnostic studies including MRI and CSF AD biomarker testing. Diagnosis of AD was made by subspecialty certified Cognitive and Behavioral Neurologists with additional input from Neuropsychologists based on current NIA-AA criteria (27,28). CSF was collected by lumbar puncture and banked according to best practice guidelines outlined by the National Institute on Aging for Alzheimer’s Disease Centers (https://alz.washington.edu/BiospecimenTaskForce.html), and identical pre-analytic steps were followed in all groups. Measurements of Amyloid-beta_1-42_ (Aβ_42_), total Tau (tTau), and phosphorylated Tau_181_ (pTau_181_) was performed on the Roche Diagnostics Elecsys platform (29-31) using recommended protocols. In total, the cohort was comprised of 105 healthy Controls and 98 AD. The racial background of each case was based upon self-identification. Of the 203 cases, 100 identified as Caucasian or White while 103 identified as African American or Black. Case metadata can be found in **Supplemental Table 1** along with a summarized version in **Table 1**.

**Table 1.**
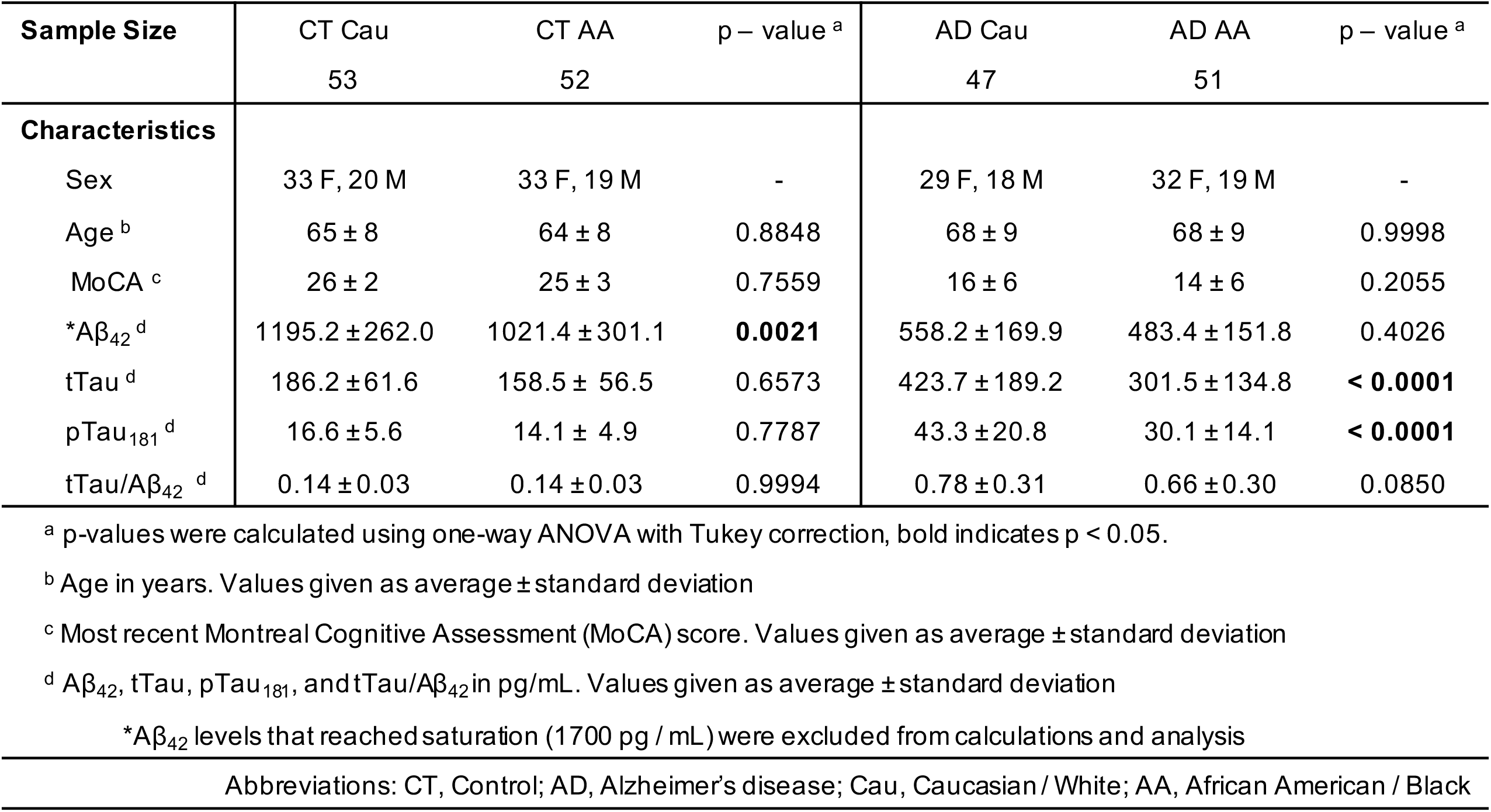
Cohort characteristics.

| Sample Size | CT Cau<br>53 | CT AA<br>52 | p – value <sup>a</sup> | AD Cau<br>47 | AD AA<br>51 | p – value <sup>a</sup> |
| --- | --- | --- | --- | --- | --- | --- |
| Characteristics |  |  |  |  |  |  |
| Sex | 33 F, 20 M | 33 F, 19 M | - | 29 F, 18 M | 32 F, 19 M | - |
| Age <sup>b</sup> | 65 ± 8 | 64 ± 8 | 0.8848 | 68 ± 9 | 68 ± 9 | 0.9998 |
| MoCA <sup>c</sup> | 26 ± 2 | 25 ± 3 | 0.7559 | 16 ± 6 | 14 ± 6 | 0.2055 |
| *Aβ<sub>42</sub> <sup>d</sup> | 1195.2 ± 262.0 | 1021.4 ± 301.1 | **0.0021** | 558.2 ± 169.9 | 483.4 ± 151.8 | 0.4026 |
| tTau <sup>d</sup> | 186.2 ± 61.6 | 158.5 ± 56.5 | 0.6573 | 423.7 ± 189.2 | 301.5 ± 134.8 | **< 0.0001** |
| pTau<sub>181</sub> <sup>d</sup> | 16.6 ± 5.6 | 14.1 ± 4.9 | 0.7787 | 43.3 ± 20.8 | 30.1 ± 14.1 | **< 0.0001** |
| tTau/Aβ<sub>42</sub> <sup>d</sup> | 0.14 ± 0.03 | 0.14 ± 0.03 | 0.9994 | 0.78 ± 0.31 | 0.66 ± 0.30 | 0.0850 |

<sup>a</sup> p-values were calculated using one-way ANOVA with Tukey correction, bold indicates p < 0.05.
<sup>b</sup> Age in years. Values given as average ± standard deviation
<sup>c</sup> Most recent Montreal Cognitive Assessment (MoCA) score. Values given as average ± standard deviation
<sup>d</sup> Aβ<sub>42</sub>, tTau, pTau<sub>181</sub>, and tTau/Aβ<sub>42</sub> in pg/mL. Values given as average ± standard deviation
\*Aβ<sub>42</sub> levels that reached saturation (1700 pg / mL) were excluded from calculations and analysis
Abbreviations: CT, Control; AD, Alzheimer’s disease; Cau, Caucasian / White; AA, African American / Black

### Protein digestion of CSF

First, 70 μL of CSF was transferred to 1 mL deep well plates for digestion with lysyl endopeptidase (LysC) and trypsin. Notably, unlike our previous study (25), these CSF samples were not immunodepleted prior to digestion. Briefly, the samples were reduced and alkylated with 1.4 μL of 0.5 M tris-2(-carboxyethyl)-phosphine (ThermoFisher) and 7 μL of 0.4 M chloroacetamide in a 90°C water bath for 10 min. The water bath was then turned off and allowed to cool to room temperature along with samples for 5 minutes. Following this, water bath sonication was performed for 5 min. The samples were then allowed to cool again to room temperature for 5 mins prior to adding urea. Then 78 μL of 8M urea buffer (8M urea, 10mM Tris, 100mM NaH_2_PO_4_, pH 8.5) and 3.5 μg of LysC (Wako), was added to each sample, resulting in a final urea concentration of 4M. The samples were then mixed well, gently spun down, and incubated overnight at 25°C for digestion with LysC. The following day, samples were diluted to 1M urea with a blend of 468 μL of 50 mM ammonium bicarbonate (32) and 7 μg of Trypsin (ThermoFisher). The samples were subsequently incubated overnight at 25°C for digestion with trypsin. The next day, the digested peptides were acidified to a final concentration of 1% formic acid and 0.1% trifluoroacetic acid. This was immediately followed by desalting on 30 mg HLB columns (Waters) and then eluted with 1 mL of 50% acetonitrile (ACN) as previously described (33). To normalize protein quantification across batches (33), 100 μl was taken from all CSF samples and then combined to generate a pooled sample. This pooled sample was then divided into global internal standards (GIS) (34). All individual samples and the pooled standards were then dried using a speed vacuum (Labconco).

### Tandem mass tag labeling of CSF peptides

All CSF samples were balanced for diagnosis, race, age, and sex (in that order) across 16 batches using ARTS (automated randomization of multiple traits for study design) (35). Using a 16-plex Tandem Mass Tag (TMTpro) kit (Thermo Fisher Scientific, A44520, Lot number: VH3111511), 13 channels of each batch were allocated to a CSF sample (127N, 127C, 128N, 128C, 129N, 129C, 130N, 130C, 131N, 131C, 132N, 132C, 133N). The remaining 3 channels were occupied with a GIS pool (126), a standard biomarker negative pool (133C), and a standard biomarker positive pool sample (134N). Information regarding the origination of these pooled samples were reported previously (36). **Supplemental Table 2** provides the sample to batch arrangement. In preparation for labeling, each CSF peptide digest was resuspended in 75 μl of 100 mM triethylammonium bicarbonate (TEAB) buffer meanwhile 5 mg of TMT reagent was dissolved into 200 μl of ACN. Once, both were in suspension, 15 μl of TMT reagent solution was subsequently added to the resuspended CSF peptide digest. After 1 hour, the reaction was quenched with 4 μl of 5% hydroxylamine (Thermo Fisher Scientific, 90115) for 15 min. Then, the peptide solutions were combined according to the batch arrangement. Finally, each TMT batch was desalted with 60 mg HLB columns (Waters) and dried via speed vacuum (Labconco).

### High-pH peptide fractionation

Dried samples were re-suspended in high pH loading buffer (0.07% vol/vol NH4OH, 0.045% vol/vol FA, 2% vol/vol ACN) and loaded onto a Water’s BEH column (2.1 mm x 150 mm with 1.7 µm particles). A Vanquish UPLC system (ThermoFisher Scientific) was used to carry out the fractionation. Solvent A consisted of 0.0175% (vol/vol) NH4OH, 0.01125% (vol/vol) FA, and 2% (vol/vol) ACN; solvent B consisted of 0.0175% (vol/vol) NH_4_OH, 0.01125% (vol/vol) FA, and 90% (vol/vol) ACN. The sample elution was performed over a 25 min gradient with a flow rate of 0.6 mL/min with a gradient from 0 to 50% solvent B. A total of 96 individual equal volume fractions were collected across the gradient. Fractions were concatenated to 48 fractions and dried to completeness using vacuum centrifugation.

### Mass spectrometry analysis and data acquisition

All samples (∼1µg for each fraction) were loaded and eluted by an Easy-nLC 1200 (Thermofisher Scientific) with an in-house packed 15 cm, 150 μm i.d. capillary column with 1.7 μm CSH (Water’s) over a 35 min gradient. Mass spectrometry was performed with a high-field asymmetric waveform ion mobility spectrometry (FAIMS) Pro front-end equipped Orbitrap Lumos (Thermo) in positive ion mode using data-dependent acquisition with 1 second top speed cycles for each FAIMS compensative voltage (CV). Each cycle consisted of one full MS scan followed by as many MS/MS events that could fit within the given 1 second cycle time limit. MS scans were collected at a resolution of 120,000 (410-1600 m/z range, 4×10^5 AGC, 50 ms maximum ion injection time, FAIMS CV of -45 and -65). Only precursors with charge states between 2+ and 5+ were selected for MS/MS. All higher energy collision-induced dissociation (HCD) MS/MS spectra were acquired at a resolution of 50,000 (0.7 m/z isolation width, 35% collision energy, 1×10^5 AGC target, 86 ms maximum ion time). Dynamic exclusion was set to exclude previously sequenced peaks for 30 seconds within a 10-ppm isolation window.

### Database search and protein quantification

All raw files were analyzed using the Proteome Discoverer Suite (v.2.4.1.15, Thermo Fisher Scientific). MS/MS spectra were searched against the UniProtKB human proteome database (downloaded in 2019 with 20338 total sequences). The Sequest HT search engine was used to search the RAW files, with search parameters specified as follows: fully tryptic specificity, maximum of two missed cleavages, minimum peptide length of six, fixed modifications for TMTPro tags on lysine residues and peptide N-termini (+304.307 Da) and carbamidomethylation of cysteine residues (+57.02146 Da), variable modifications for oxidation of methionine residues (+15.99492 Da), serine, threonine and tyrosine phosphorylation (+79.966 Da) and deamidation of asparagine and glutamine (+0.984 Da), precursor mass tolerance of 10 ppm and a fragment mass tolerance of 0.05 Da. Percolator was used to filter peptide spectral matches and peptides to an FDR <1%. Following spectral assignment, peptides were assembled into proteins and were further filtered based on the combined probabilities of their constituent peptides to a final FDR of 1%. Peptides were grouped into proteins following strict parsimony principles. A complete TMT reporter ion abundance-based table output of assembled protein abundances without adjustments can be found in **Supplemental Table 3**.

### Adjustment for batch and other sources of variance

Only proteins quantified in ≥ 50% of samples were included in subsequent analysis (n = 1840 proteins). As previously reported (23-25), batch correction was performed using a Tunable Approach for Median Polish of Ratio, (https://github.com/edammer/TAMPOR), an iterative median polish algorithm for removing technical variance across batch. Multidimensional scaling (MDS) plots were used to visualize batch contributions to variation before and after batch correction Noticeably, prior to batch correction, cases within the same batch clustered together and batches ran consecutively tended to cluster more closely together (**Supplemental Figure 1A)**. Following batch correction using a median polish algorithm, the cases were no longer clustering by batch (**Supplemental Figure 1B**). The data was then subjected to outlier removal using a robust principal component analysis method, *PcaGrid* (37). A scree plot graphing the eigenvalue against the principal component (PC) number was utilized to determine the number of PCs to include in the parameters (**Supplemental Figure 1C**). Briefly, the parameters used for outlier detection were as follows: the desired number of principal components = 7, method = mean absolute deviation, and criterion for computing cutoff values = 0.99 (**Supplemental Figure 1D**). This resulted in the detection and removal of 15 outliers, resulting in a final n=189 samples. Bootstrap regression was then performed to remove for covariates such as age at collection and sex. Variance partition analysis was performed to confirm appropriate regression of these traits (**Supplemental Figure 1E & 1F**). Since the *variancePartition* package does not allow missing values, proteins with missing quantifications were temporarily imputed using the *impute*.*knn* function of the impute R package. The final cleaned dataset after regression and log2 transformation can be found in **Supplemental Table 4**.

### Differential expression analysis

One-way ANOVA followed by Tukey’s post hoc adjustment for multiple comparisons was performed on four groups (Control-Caucasian, Control-African American, AD-Caucasian, and AD-African American) to identify differentially expressed proteins across diagnosis and within each race. Differentially expressed proteins for comparisons of interest (i.e., Control-Caucasian vs AD-Caucasian and Control-African American vs AD-African American) were then presented as volcano plots using the *ggplot2* package in R v4.1.2. A list of all comparisons computed with corresponding adjusted p-values is provided in **Supplemental Table 5**.

### Weighted Gene Co-expression Network Analysis (WGCNA)

As previously published (21,23-25), the blockwiseModules function from the WGCNA package in R was utilized to derive the weighted protein co-expression network (**Supplemental Table 6**). Briefly, the following parameters were used: soft threshold power beta = 3, deepSplit = 4, minimum module size = 5, merge cut height = 0.07, and a signed network with partitioning about medoids. Using the *pairwise*.*wilcox*.*test* R function with Bonferroni correction, a pairwise Wilcox test was performed to calculate pairwise comparisons between each group with corrections for multiple testing.

### Gene ontology (GO) and cell type enrichment analysis

To characterize co-expressed protein module biology, gene ontology (GO) annotations were retrieved from the Bader Lab’s monthly updated .GMT formatted ontology lists downloaded July 5, 2022 (38). A Fisher’s exact test for enrichment was performed into each module’s protein membership using an in-house script (https://github.com/edammer/GOparallel). Significant ontologies for each module are included in **Supplemental Table 7**. For cell type enrichment analysis, an in-house marker list was as previously described (23) (**Supplemental Table 8**). A Fisher’s exact test was performed for each module member list using the merged human cell type marker list to determine cell type enrichment. For brain-CSF module overlap a one-sided Fisher’s exact test to compare significance of module membership.

### Selected Reaction Monitoring

Selected reaction monitoring (SRM) assays were performed on 195 of the 203 cases to determine whether a separate targeted proteomic approach could replicate proteomic changes seen in TMT discovery proteomics. Sample preparation, peptide quantification, and data acquisition for the SRM assay was performed as previously described (36). Briefly, peptides were selected based on their robust detection and significant differential expression in previous CSF discovery proteomic projects for synthesis as heavy standards (25,39). In addition to the 195 samples from before, two pools of CSF were utilized as AD biomarker positive and AD-biomarker negative quality controls standards (36). After all of the CSF samples were blinded and randomized, each sample (50 μL) was reduced, alkylated, denatured and then subjected to digestion as described (36). After digestion, the heavy labeled standards, 15uL per 50 μL of CSF, were added to the peptides. The peptides then underwent acidification, desalting and were dried under vacuum. The peptides were quantified using TSQ Altis Triple Quadrupole mass spectrometer as described (36). The resulting raw files were uploaded to Skyline-daily software (version 21.2.1.455) for peak integration and quantification by peptide ratios. The technical coefficient of variation (CV) of each peptide was then calculated based on the peptide area ratio for the AD-positive and AD-negative quality control pools (**Supplemental Table 9**). CSF peptide targets with CVs ≤ 20% in at least one pooled standard were determined as peptides with high precision and were kept for subsequent analysis (n = 85 peptides). The peptide ratios for these remaining peptides were then log_2_ transformed and true zero values were replaced after log_2_-transformation with the minimum value for that peptide minus 1 (**Supplemental Table 10)**. Bootstrap regression was then used to regress for age and sex (**Supplemental Table 11**). Bicor was then utilized to calculate the correlation between SRM peptides and TMT-MS protein measurements (**Supplemental Table 12**). In cases where multiple peptides mapped to one protein, the most correlated peptide was kept for further analysis (**Supplemental Table 13**). One-way ANOVA analysis with Tukey adjustment was then utilized once again to examine pairwise interactions **(Supplemental Table 14**) and receiver operating characteristic (ROC) curve analysis was performed as previously described (36) (**Supplemental Figure 15**).

### Data and Code Availability

Raw mass spectrometry data and pre- and post-processed protein expression data and case traits related to this manuscript are available from CSF proteomics can be found at https://www.synapse.org/EmoryDiversityCSF. The results published here are in whole or in part based on data obtained from the AMP-AD Knowledge Portal (https://adknowledgeportal.synapse.org). The AMP-AD Knowledge Portal is a platform for accessing data, analyses and tools generated by the AMP-AD Target Discovery Program and other programs supported by the National Institute on Aging to enable open-science practices and accelerate translational learning. The data, analyses and tools are shared early in the research cycle without a publication embargo on secondary use. Data are available for general research use according to the following requirements for data access and data attribution (https://adknowledgeportal.synapse.org/#/DataAccess/Instructions).

## RESULTS

### Cohort characteristics

The main objective of this study was to perform an unbiased proteomic study of CSF to define biomarkers and pathways that are similar or divergent between African Americans and Caucasians with AD. **Figure 1A** provides an overview of our study approach, which included the generation of balanced sets of CSF samples from African American and Caucasian individuals, matched for age and sex with roughly equal numbers of controls and AD cases (**Table 1**). This included 53 Caucasian controls, 52 African American controls, 47 AD Caucasians, and 51 AD African Americans. The majority were female and on average the controls (64.5 years) were slightly younger than AD (68 years). Notably, there were no statistical differences between the ages of the African Americans and the Caucasians within diagnosis (control: p=0.8848, AD: p=0.9998). As expected, AD cases had lower Montreal Cognitive Assessment (MoCA) scores than controls, but there were no statistically significant differences between MoCA scores across race within controls and AD (control: p=0.7559, AD:p=0.2055). The AD cases also had lower Amyloid-beta (Aβ_42_) levels and elevated total Tau (tTau) and phosphorylated Tau_181_ (pTau_181_) levels. Notably, Aβ_42_ levels were significantly lower in African American Controls compared to Caucasian controls (p = 0.0021) but not different between African American AD and Caucasian AD. Conversely, tTau and pTau_181_ levels were significantly lower in African Americans with AD (tTau:p<0.0001, pTau_181_:p<0.0001) but not different between African American and Caucasian controls.

**Figure 1.**
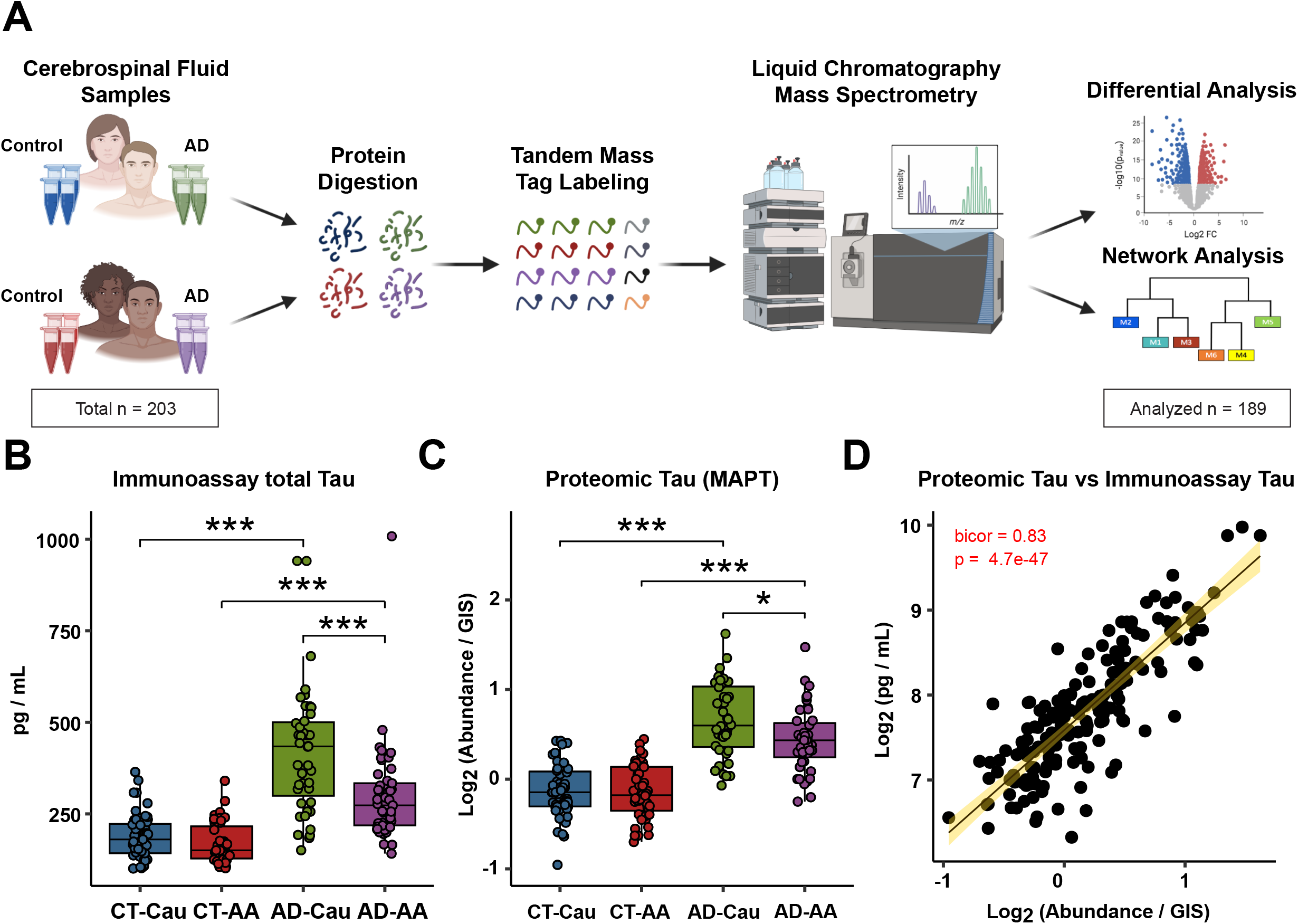
Schematic of experimental workflow and correlation between proteomic Tau and total Tau immunoassay measurements. (**A**) Schematic of experimental workflow for quantification of cerebrospinal fluid proteome. (**B**) Total Tau levels as measured by Roche Elecsys Platform between control (CT) and AD cases and stratified by self-identified race: Caucasian (Cau) or African American (AA) (**C**) Tau levels measured by mass spectrometry. One-way ANOVA with Tukey post-hoc correction determined pairwise relationships (**D**) Correlation of Tau levels by TMT-MS (x-axis) to paired immunoassay total Tau levels (y-axis). Biweight midcorrelation coefficient (bicor) with associated p-value is shown. Only 179 cases were included in the linear regression analysis because of sample outlier removal and missing values in the TMT-MS.

### Mass spectrometry and immunoassay measurements of Tau strongly correlate

Following enzymatic digestion, TMT labeling, and off-line fractionation, all samples were subjected to liquid chromatography tandem mass spectrometry (LC-MS/MS) (**Figure 1A**). In total, TMT-MS proteomic analysis identified 34,330 peptides mapping to 2,941 protein groups across the 203 samples (16 total batches). To account for missing protein measurements across batches, we included only those proteins quantified in at least 50% of samples following outlier removal as previously described (21-25), resulting in the final quantification of 1,840 proteins. Protein abundance was adjusted for batch and age and sex were regressed. Protein levels of Tau (MAPT) by TMT-MS correlated strongly to independently measured tTau levels via immunoassay (r=0.83, p = 4.7e-47). As expected, Tau levels were significantly elevated in both African Americans and Caucasians with AD across both platforms compared to controls (**Figure 1B and 1C**). Consistent with the immunoassay measurements, TMT-MS Tau levels also showed significantly lower levels in African Americans with AD compared to Caucasians with AD (**Figure 1C**). Thus, in this study, both platform measures of CSF Tau support a reduction of total Tau levels in African Americans with AD, consistent with previous findings (16,17).

### Differential expression analysis of African American and Caucasian CSF proteome reveals unique and shared changes in AD

Differential expression analysis was performed to identify changes in the CSF proteome by race in AD (**Supplemental Table 5**). Consistent with previous proteomic analyses of AD CSF (24,25,40-42), there was a significant increase in Tau (MAPT), 14-3-3 proteins, (YWHAZ, YWHAG, and YWHAE), SMOC1, neurofilaments (NEFM and NEFL) and proteins involved in glucose metabolism in both African Americans and Caucasians with AD compared with race matched controls (**Figure 2A & 2B**). However, Caucasians with AD exhibited a bias towards proteins that were increased in AD, where the number of differentially expressed proteins (DEPs) was nearly double (n=183 proteins) the number of decreased DEPs in AD (n=74 proteins) (**Figure 2A**). In contrast, in African Americans the number of increased and decreased DEPs was more balanced (151 increased proteins vs. 162 decreased proteins). A Venn diagram illustrates the overlap of DEPs from African Americans and Caucasians with AD (**Figure 2C**), with the majority of proteins (n=168 proteins) differentially expressed in both races. Furthermore, a correlation analysis of both shared and unique DEPs showed overall high agreement in direction of change (bicor=0.887, p=2.47e-136, **Figure 2D)**. However, there were some exceptions including SLIT1 and VSTM2A, which were significantly increased in Caucasians, but decreased in African Americans with AD. Both proteins are predominantly enriched in neuronal-cell types (43,44). Thus, despite the differences in the number of significant DEPs in African Americans compared to Caucasians with AD, the direction of change with disease remains largely similar across both races.

**Figure 2.**
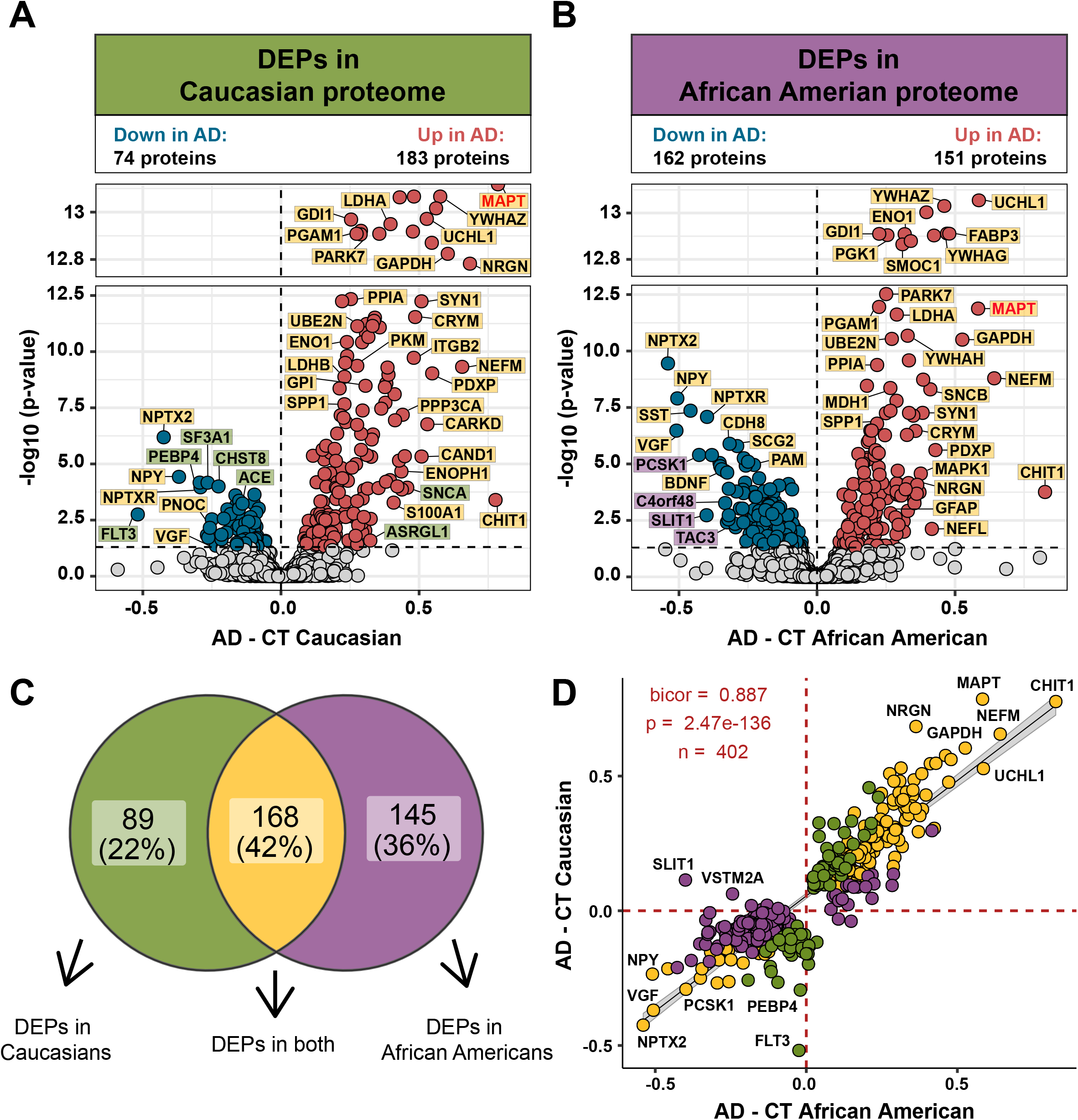
Differential expression of Caucasian and African American CSF proteomes in AD. Volcano plot displaying the log_2_ fold change (FC) (x-axis) against one-way ANOVA with Tukey correction derived -log10 p-value (y-axis) for all proteins (n=1840) comparing AD versus Controls for Caucasians (**A**) and African Americans (**B**). Cutoffs were determined by significant differential expression (p<0.05) between control (CT) and AD cases. Proteins with significantly decreased levels in AD are shown in blue while proteins with significantly increased levels in disease were indicated in red. Select proteins were denoted and labeled by whether they were differentially expressed in both proteomes (yellow), in only the Caucasian proteome (green), or in only the African American proteome (purple). (**C**) Venn diagram illustrating the number of differentially expressed proteins (DEPs) that were uniquely changed in one proteome (green or purple) or changed in both proteomes (yellow) (**D**) The correlation between the fold change of all DEPs (n=402) across the African American proteome (x-axis) and the Caucasian proteome (y-axis) were strongly correlated (bicor=0.887, p=2.47e-136), regardless of whether the DEP was significant in one (green or purple) or both proteomes (yellow).

### Network analysis of the CSF proteome reveals modules related to pathways and brain cell-types

Co-expression network analysis of the AD brain proteome organizes proteins into modules related to molecular pathways, organelles, and cell types impacted by AD pathology (21-24). Moreover, integration of the human AD brain and CSF proteome revealed that approximately 70% of the CSF proteome overlapped with the brain proteome (25). While proteomic networks in AD brain have been examined, network changes in the AD CSF proteome, including those associated with race and AD biomarkers are less well understood. Thus, we applied Weighted Gene Co-expression Network Analysis (WGCNA) to define trends in protein co-expression across 1840 CSF proteins in all individuals. These parameters identified 14 modules (M), ranked by size, ranging from the largest M1, with 370 proteins to the smallest, M14, with 16 proteins (**Figure 3A**). Many of these modules were significantly enriched for brain-specific cell types (**Figure 3B**) as well as established brain gene ontologies (GO), cellular functions and/or organelles (**Figure 3C, Supplemental Table 7**). The three largest modules were associated with categories of “Postsynaptic Membrane” (M1), “Complement Activation” (M2), and “Extracellular Matrix” (M3) whereas M5 represented “Lysosome” and M6 “Gluconeogenesis”. Other modules included those with GO terms linked to “Cell Morphogenesis” (M4), “Cell Redox / Homeostasis” (M7), “Protein Ubiquitination” (M8), “Angiogenesis / Epithelial Cell Migration” (M9), “Synapse Assembly / Lysosome” (M10), Myofibril Assembly (M11), “Actin Cytoskeleton” (M12), “Kinase Signaling” (M13), and “Carbohydrate Metabolism” (M14).

**Figure 3.**
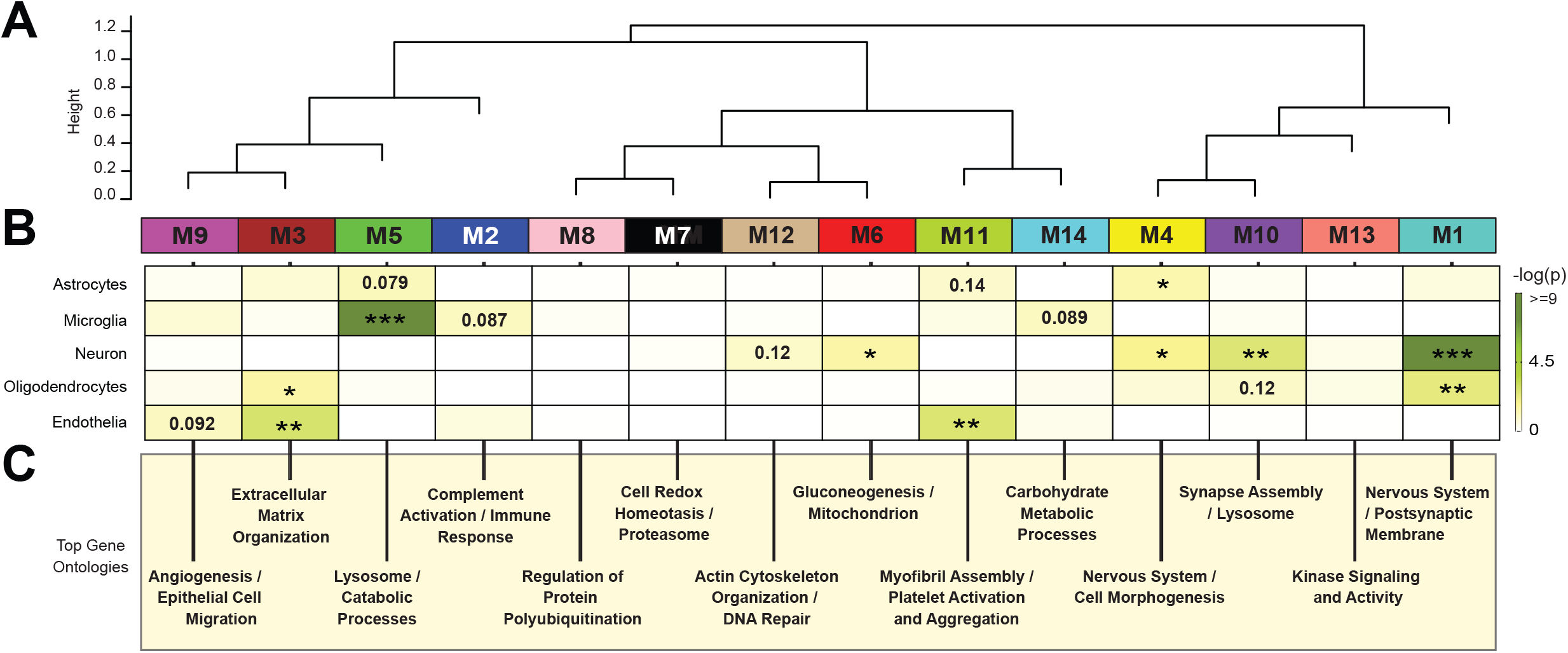
Network analysis classifies the CSF proteome into modules associated with specific brain cell-types and gene ontologies. (**A**) Weighted Gene Co-expression Network Analysis cluster dendrogram groups proteins (n=1840) into 14 distinct protein modules (M1-M14). (**B**) Cell-type enrichment was assessed by cross referencing module proteins by matching gene symbols using a one-tailed Fisher’s exact test against a list of proteins determined to be enriched in neurons, oligodendrocytes, astrocytes, microglia and endothelia. The degree of cell-type enrichment increases from yellow to dark green with asterisks denoting the following statistical significance (*p≤0.05; **p≤0.01; ***p≤.001). Top gene ontology (GO) terms were selected from significant GO annotations.

Protein-based network analysis in AD brain tissue has shown that the cellular composition represents a major source of biological variance and that many of the network modules are enriched in proteins that are expressed by specific brain cell types (23,24). To determine if a similar relationship exists with protein-based networks in CSF, we evaluated the overlap of proteins in each module with brain cell-type specific makers (**Figure 3B, Supplemental Table 8**), generated previously from cultured or acute isolated neurons, oligodendrocytes, astrocytes, endothelial, and microglia from brain (43,44). The largest module M1, was enriched with neuron/synaptic proteins (i.e., NPTX1, NPTXR, SCG2, VGF, NRN1, and L1CAM) and to a lesser degree oligodendrocyte proteins (i.e., IGSF8, VCAN, APLP1). The M4 module was also enriched for neuronal protein markers including RTN4R1, LINGO2, OLFM1, and PLXNA2, associated with “Nervous Systems and Cell Morphogenesis”. Modules most enriched with microglia markers were M2 (i.e., C2, C3, C1RL, C1QA, C1QB, C1QC, LCP1, etc.) and M5 (i.e., HEXB, CTSZ, HEXA, CTSA, CTSB) consistent with a role in complement activation and lysosome function, respectively. Finally, endothelial markers were mainly overrepresented in modules M3 (i.e., NID2, ECM2, NID1, LTBP4, LAMA5, LAMC1), M9 (IGFBP7, F5, SDCBP, BGN) and M11 (FLNA, ANXA5, S100A11, MYL6) consistent with roles in extracellular matrix, angiogenesis and myofibril assembly, respectively. Thus, as seen in the network analysis of bulk proteome from human brain (23,24), certain modules of co-expressed proteins in CSF were enriched with markers of specific brain cell-types. To further support this observation, we assessed the protein overlap between modules in CSF and modules from a recent large-scale consensus TMT-MS proteomic network of bulk human AD brain tissue (23). (**Supplemental Figure 2**). Except for M9, M10 and M14, which had minimal overlap with the brain, all other modules (79% total) in the CSF network significantly overlapped with at least one of the 44 brain modules (B-M1 to B-M44). For example, there is overlap with proteins in M1 “Postsynaptic Membrane” with several neuronal modules in the consensus brain network (B-M1, B-M4, B-M5, B-M10, and B-M15). In addition, M2 “Complement Activation” in CSF overlaps with modules in human brain associated with complement and immune response (B-M26 and B-M40), whereas M3 “Extracellular Matrix” strongly overlap with B-M27 in brain enriched with endothelial cell markers (**Supplemental Figure 2**). Collectively, this supports that the co-expression in protein levels is, in part, shared between CSF and brain tissue, which could reflect changes in activation or phenotypes of specific brain cell types.

### CSF protein modules correlate to race and clinicopathological phenotypes of AD

We assessed module correlation to race, cognitive scores (MoCA), and the hallmark AD biomarkers Aβ_42_, tTau, and pTau_181_. The protein network resulted in three main groups/clusters based on module relatedness (**Figure 4A)**. The first group (Group 1) was comprised of four modules (M2 “Complement”, M5 “Lysosome/Microglia”, M3 “Extracellular Matrix”, and M9 “Angiogenesis / Cell Migration” that exhibited baseline racial differences in abundance levels, irrespective of disease diagnosis (**Figure 4B**). Notably, the eigenprotein, which corresponds to the first principal component of a given module and serves as a summary expression profile for all proteins within a module, were increased for all four modules in African Americans compared to Caucasians. Of note, these modules are significantly enriched with microglia, astrocyte (M2 and M5) and endothelial cell markers (M3 and M9), which suggests that genetic ancestry and/or environmental differences influence expression or secretion of these cell-type markers.

**Figure 4.**
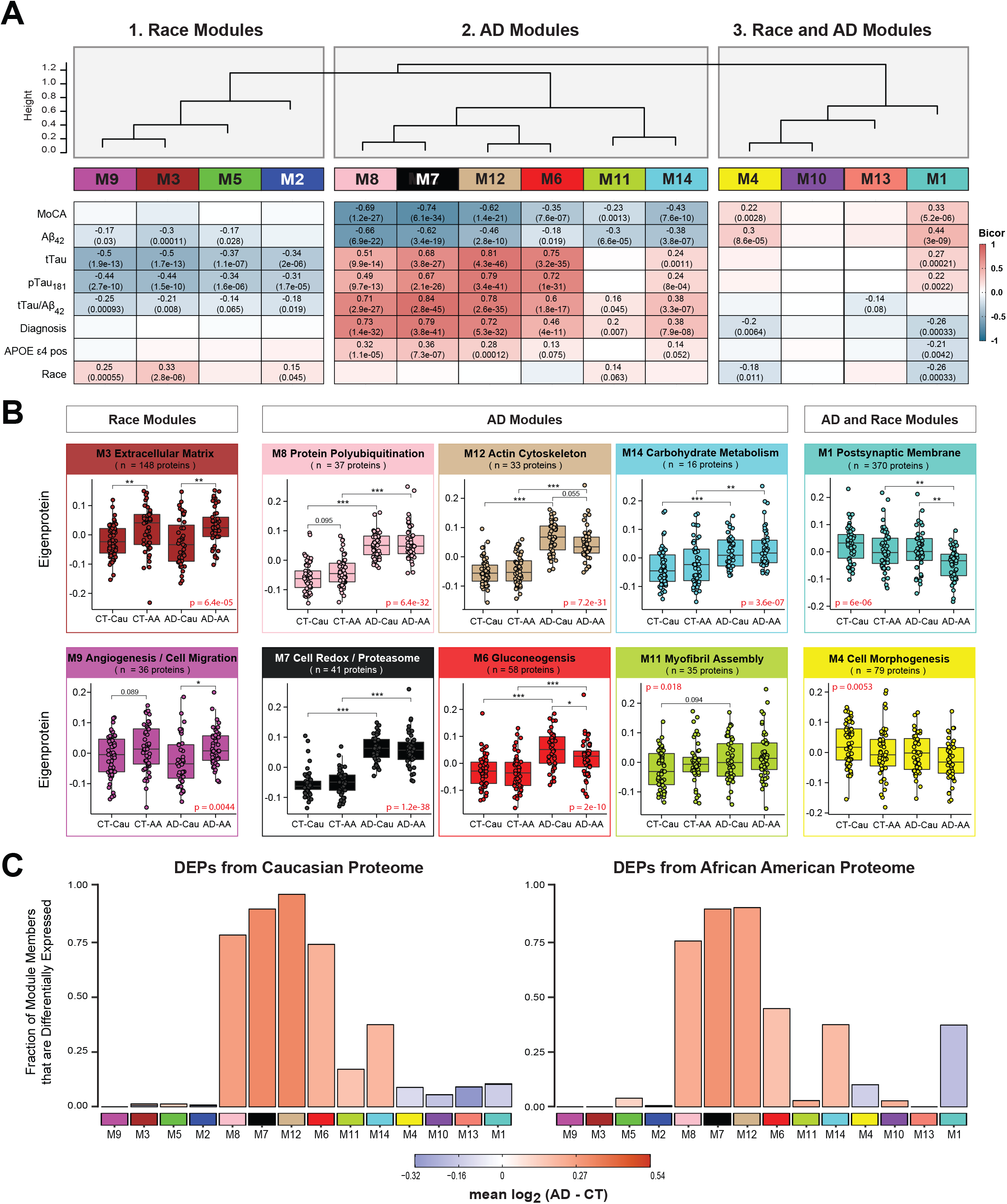
CSF protein modules correlate to race and clinicopathological phenotypes of AD. (**A**) Modules were clustered based on relatedness defined by correlation of protein co-expression eigenproteins (indicated by position in color bar). There were three main clusters in the network: Race-specific (Group 1), AD-specific (Group 2), and AD- and race-specific modules (Group 3). Biweight midcorrelation (bicor) analysis of module eigenprotein levels with diagnostic measures of AD, including MoCA score, immunoassay Amyloid-beta_1-42_ (Aβ_42_), total Tau (tTau), phosphorylated Tau_181_ (pTau_181_), ratio measures of tTau/Aβ_42_, diagnosis, whether the sample has *APOE* ε4 allele or not, and race. The strength of positive (red) and negative correlations are shown by a heatmap with annotated bicor correlations and associated p-values. (**B**) Eigenprotein values distributed by race and diagnosis of representative modules for each cluster. (**C**) Differential protein abundance AD samples compared to controls, by module with Caucasian proteome on the left and African Americans on the right. The height of the bars represents the fraction of module member proteins that DEPs compared to controls. The bars are color coded by heatmap for average log_2_ difference in abundance, where red represents an increase in abundance in AD, and blue represents a decrease in abundance in AD.

The second cluster of modules (Group 2) was comprised of six modules that were all increased in AD (M14, M11, M6, M12, M7, and M8) (**Figure 4A**). These AD modules also demonstrated significant negative correlations to MoCA scores and, conversely, significant positive correlations to tTau/Aβ_42_ ratio and APOE ε4 risk (**Figure 4A**). Interestingly, a hub protein of M12 Actin/Cytoskeleton module was Tau (MAPT). Consistent with CSF levels observed for Tau by immunoassay and TMT-MS (**Figure 1**), the M12 eigenprotein had lower levels in African Americans, compared to Caucasians with AD, albeit not significant (p=0.055) (**Figure 4B**). Notably, M6 Gluconeogenesis was significantly lower in African Americans compared to Caucasians with AD, highlighting another module of CSF proteins that differed between race in AD (**Figure 4 A and B**). This also indicates that the increased glycolytic signature of AD previously reported in CSF (24,25) is higher in Caucasians with AD. Consistently, a greater proportion of increased DEPs in Caucasians with AD mapped to M6 compared to African Americans with AD (**Figure 4C)**. In contrast, M7 “Cell Redox/ Homeostasis” and M8 “Protein Ubiquitination”, had the strongest correlations to tTau/Aβ_42_ ratio and cognition (**Figure 4B**), yet both demonstrated strong, equivalent elevations in African Americans and Caucasians with AD (**Figure 4B**). This is consistent with an equivalent fraction of increased DEPs mapping to these modules in African American and Caucasians with AD (**Figure 4C)**. Therefore, proteins in these modules including 14-3-3 family members (YWHAZ, YWAHB, YWHAG, YWHAE) likely represent the best class of CSF AD biomarkers that are not influenced by race. M14 Carbohydrate Metabolism and M11 Myofibril Assembly were both elevated in both African Americans and Caucasians with AD (**Figure 4A and B**), yet to a lesser degree than M7 and M8.

The final group of modules (Group 3) contained neuronal modules M1 “Postsynaptic Membrane” and M4 “Nervous System/Cell Morphogenesis” that showed strong correlation to both race and AD diagnosis (**Figure 4A**). Both modules are i) decreased in African Americans compared to Caucasians and ii) decreased in AD compared to controls. M1 and M4 are also enriched with neuronal markers and positively associated with cognitive MoCA scores (**Figure 4A**). Pairwise statistical analysis for eigenprotein levels of M1 across diagnosis and race revealed significantly lower levels in African Americans with AD (**Figure 4B**). To this end, most of the decreased DEPs in African Americans with AD mapped to M1 and to a lesser degree M4, whereas decreased DEPs in Caucasians with AD were equally distributed to M1, M4, M13 and M10 (**Figure 4C**). Notably, M10 and M13 within Group 3 did not show any differences with AD or race and did not significantly correlate with traits explored in this study (**Figure 4 and Supplemental Figure 3**). Overall, network analysis effectively organizes the CSF proteome into protein modules that are strongly linked to hallmark AD biomarkers (Aβ_42_, tTau and pTau_181_) and cognition, which in some cases were also influenced by race.

### Validation of Shared and Divergent CSF Protein Alterations across AD and Race

To further validate these network findings, we used a targeted mass spectrometry method, selected reaction monitoring (SRM), with heavy labeled internal standards to measure CSF proteins across 195 of the 203 cases included in the discovery TMT-MS assays (**Figure 5A**). The proteins and corresponding targeted peptides were previously selected based on their robust detection and significant differential expression in previous CSF discovery proteomic datasets (25,39). We used pooled CSF samples of control, and AD cases as quality controls replicates (n=29 samples total) to assess technical reproducibility. Of the peptides targeted, 85 (mapping to 58 proteins) had a coefficient of variation of <20% in both the control and AD pools with no missing values (**Supplemental Table 9 and 10**). Following adjustments of co-variates (i.e., age and sex), peptide levels were highly correlated with protein levels measured by TMT-MS from the same samples (**Supplemental Tables 11 and 12**). If a protein was measured by more than one peptide the most correlated peptide to the TMT-MS protein level was selected for further analysis. The final peptide list can be found in **Supplemental Table 13**. ANOVA analyses determined pairwise significance between the four groups (i.e., Control-Caucasians vs Control-African Americans vs AD-Caucasians vs AD-African Americans, **Supplemental Table 14**). **Figure 5B** highlights peptides (n=24) that reached significance and that mapped to proteins in CSF modules associated with race and/or AD. Consistent with the TMT-MS protein measurements, proteins measured by SRM within M7 (GAPDH and YWHAG) and M8 (YWAHB and PPIA) had strong elevations (p < 0.001) in abundance in AD in both races, whereas proteins in M12 (SMOC1, PARK7, and LDHB) had a greater magnitude of change in Caucasians than African Americans with AD. Similarly, a majority of the proteins measured by SRM in M6 (PKM, GDA, TPI1, GOT1, ALDOA and ENO2) were more increased in Caucasians than African Americans with AD (**Figure 5B and 5C**). Proteins in the synaptic M1 module (VGF, SCG2, NPTX2, and NPTXR) were significantly decreased in African Americans with AD compared to Caucasians (**Figure 5B and 5C**), again consistent with TMT-MS protein level abundance. Notably, African Americans with or without APOE ε4 allele in the AD group had reduced levels of these CSF peptide biomarkers compared to Caucasians indicating that race and not APOE status was driving the difference in abundance (**Supplemental Figure 4**). Finally, a receiver operating characteristic (ROC) curve analysis was performed to assess the performance of each peptide biomarker in differentiating controls and AD by race (**Figure 6 and Supplemental Table 15**). We generated an area under the curve (AUC) for AD in African American and Caucasian individuals for each protein biomarker (considered separately in each race). As expected, proteins mapping to M8 and M7 including 14-3-3 proteins (YWHAB, YWHAG and YWHAZ) were equally able to discriminate AD from control irrespective of racial background. Notably, despite having lower levels in African Americans with AD compared to Caucasians with AD, only a modest improvement in the AUC for SMOC1 was observed for classifying AD in Caucasians AUC=0.8255 (p=1.71e-08, CI=0.7421-0.9090) compared to African Americans AUC=0.7618 (p=4.12e-06, CI=0.6660-0.8576). Similar findings were observed for another M12 protein, LDHB, as well as M6 proteins PKM and ALDOA. However, the M1 protein VGF was only nominally significant at classifying AD in Caucasian AUC=0.6030 (p=0.0406, CI=0.4887-0.7173), yet highly significant in African Americans AUC=0.7593 (p=5.03e-06, CI=0.6634-0.8552). Similar results were observed for other synaptic M1 proteins, NPTX2 and SCG2, whereas NPTXR showed only a modest improvement in the AUC between African Americans compared to Caucasians with AD (**Figure 6 and Supplemental Table 15**). Collectively this supports a hypothesis that African Americans with AD have lower levels of a subset of neuronal biomarkers compared to Caucasians with AD.

**Figure 5.**
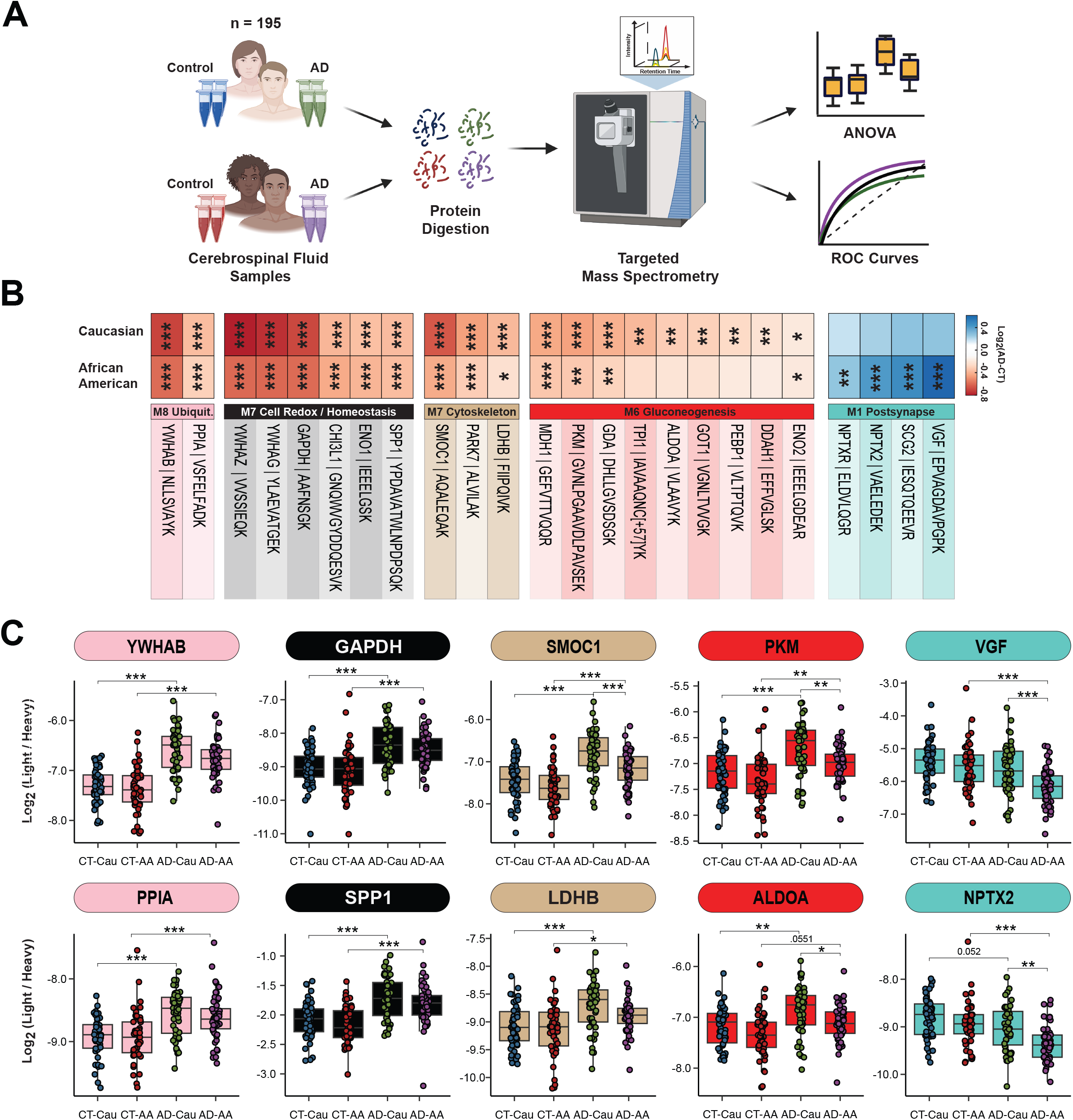
Validation of shared and divergent CSF protein levels across AD and race. (**A**) Schematic of experimental workflow for SRM analysis of cerebrospinal fluid proteome (**B**) Heatmap of peptides that were significantly differentially expressed between Control and AD Caucasians or African Americans. Stars are indicative of the level of significant difference (*p≤0.05; **p≤0.01; ***p≤0.001) seen for each peptide between AD and Control within each race. Meanwhile the colors are indicative of the log_2_ fold change (FC) of each peptide from Control and AD for each race where blue is indicative of the degree of decrease and shade of red is indicative of the degree of increase. (**C**) Log_2_ abundance of peptides that mapped to modules of interest distributed by race and diagnosis. Pairwise significance was calculated using one-way ANOVA with Tukey adjustment.

**Figure 6:**
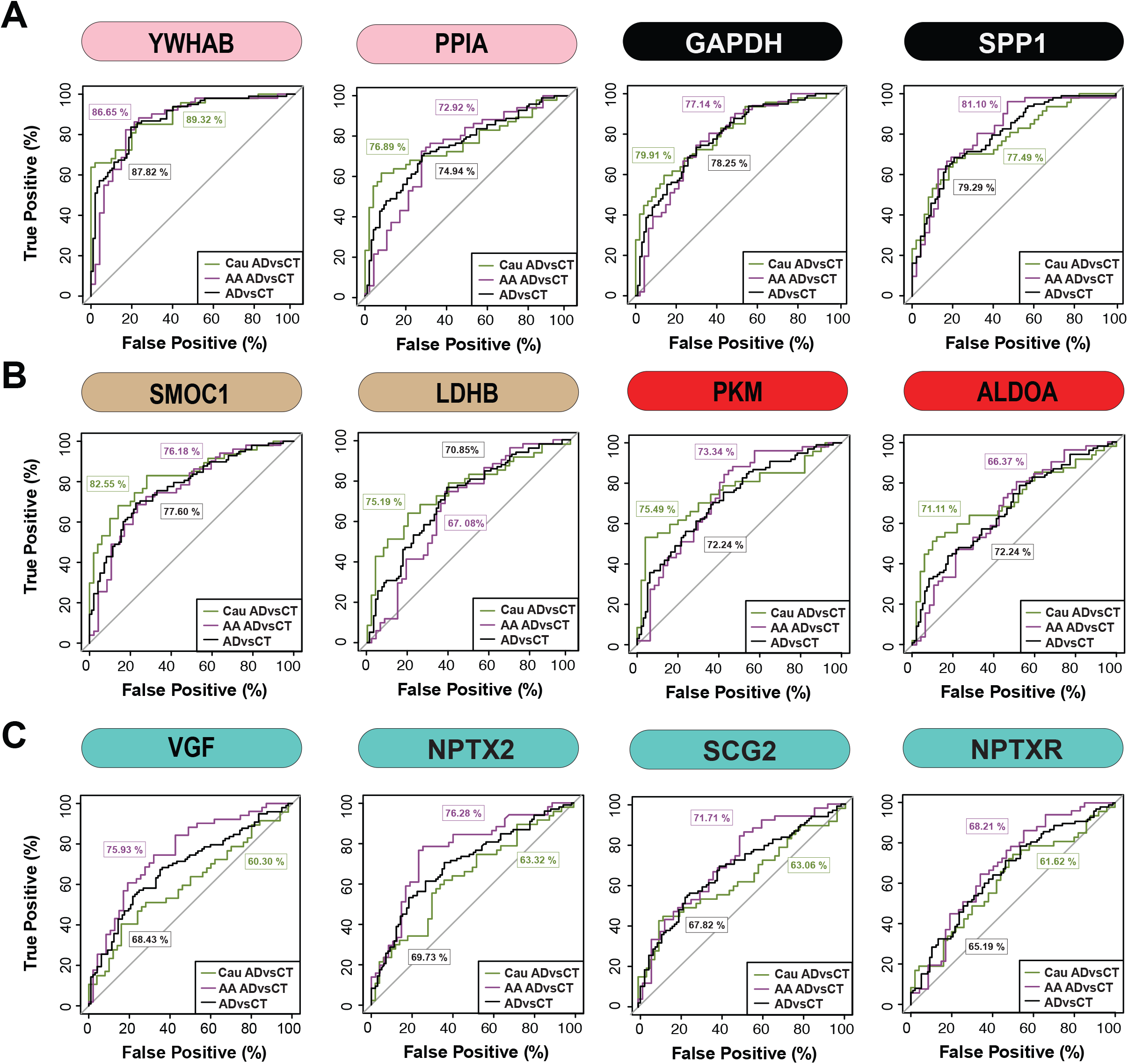
ROC analysis to evaluate CSF protein classification of AD by race. (**A**) YWHAB, PPIA, GAPDH, and SPP1 had similar performance in classifying Caucasians and African Americans with AD (**B**) SMOC1, PKM, LDHB and ALDOA showed modest improvement in the AUC for Caucasians with AD compared to African Americans with AD (**C**) VGF, SCG2, and NPTX2 were better classifiers for AD in African Americans compared to Caucasians, whereas NPTXR showed modest improvement in classification of AD in African Americans. All protein AUCs with p-values and confidence internals (CI) are provided in **Supplemental Table 15**.

## DISCUSSION

Here we performed an unbiased quantitative analysis of the AD CSF proteome to identify protein biomarkers reflective of underlying brain physiology that are shared or unique across race. Using a network analysis, we organized the CSF proteome into 14 modules of highly correlated proteins. Notably, these modules were associated with cell-types and biological pathways in brain and largely overlapped with modules in a consensus human AD brain proteomic network (23). Consistent with previous findings (16,17), we also show that Tau levels are lower in African Americans with AD compared to Caucasians. Notably, Tau mapped to a CSF module enriched with other related neuronal/cytoskeletal proteins with a magnitude of increase greater in Caucasians than in African Americans with AD. In contrast, CSF modules which included 14-3-3 proteins, were elevated equivalently in both African Americans and Caucasians with AD. A module enriched with neuronal/synaptic proteins including VGF, SCG2, and NPTX2 was significantly lower in African Americans than Caucasians with AD. VGF, SCG2, and NPTX2 levels in CSF measured by SRM were also significantly better at classifying African Americans with AD than Caucasians. Thus, our findings suggest that there are likely distinct mechanisms underlying the abundance and/or secretion of neuronal markers including Tau and VGF that differ by race. Collectively, these data underscore the need for further investigations into how AD biomarkers and underlying physiology vary across different racial backgrounds.

In a previous study we performed unbiased TMT-MS from a small discovery cohort of control and AD CSF samples (n=40) and mapped these proteins onto a human AD brain co-expression network, which revealed that approximately 70% of the CSF proteome overlapped with the brain network (25). The increased sample size in this study afforded the opportunity to build an independent co-expression network of the CSF proteome and then assess overlap with a consensus brain network. We observed a strong overlap between CSF and brain modules, indicating that there is conservation of protein co-expression across brain and CSF. This correlation is, in part, driven by the cell-types changes, as we observed modules that were enriched with neuronal, glial and endothelial specific markers. In terms of the CSF network biology that differed by race, it is noteworthy that modules significantly enriched in endothelial proteins (M3 and M9) were increased in African Americans across both control and AD individuals. This suggests that fundamental differences exist in the levels and/or activation state of cells residing the vasculature between African Americans and Caucasians. Whether this biological difference between the two races is observed in brain tissues or relates to a higher incidence of vascular health disparities between African Americans and Caucasians (45) requires further investigation.

The current biological framework for the pre-symptomatic stages of AD is based on the presence of β-amyloidosis (A), tauopathy (T), and neurodegeneration (N) also termed the A/T/N framework (46). CSF remains the gold standard for A/T/N biomarkers of neurodegenerative disease as it maintains direct contact with the brain and reflects biochemical changes amyloid, tau and neurodegeneration. A strength of our study was the balanced nature of African American samples, which offered the ability to examine racial differences in both cognitively normal controls with biomarker positive (A+/T+) individuals diagnosed with AD. Our mass spectrometry measurements of Tau strongly correlated with immunoassay levels measured on the Roche Elecsys platform reinforcing measurements made by TMT-MS. Increased Tau in CSF is considered to result from neurodegeneration, however, it can be increased in early pre-symptomatic disease stages when neurodegeneration is limited (46,47). Recently, Tau CSF levels have been linked to enhanced synaptic plasticity (48). A large proportion of synaptic proteins in this study mapped to M1 and M4 that positively associate with cognition and are decreased in AD. However, other modules, M6 and M12, also contained neuronal proteins including Tau (MAPT), SNCB, SYN1, BASP1, GAP43, SYT1, NRGN among others that were negatively associated with cognition and increased in AD. Thus, there are likely distinct mechanisms that result in the activation or abundance of neurons that contribute to this discordance in levels in AD CSF. We observed that African Americans in this study had on average lower levels of neuronal markers mapping to M1 and M4 in the network, which are reduced in AD. Paradoxically, African Americans also have lower levels of neuronal proteins in M6 and M12, which all increase in AD. Thus, increased CSF Tau, thought to be a marker of neurodegeneration, does not equate to decreased levels of VGF and other postsynaptic markers in CSF. Consistent with this observation, in a recent proteomic study of pre-symptomatic AD cases in a mainly Caucasian European population stratified by Tau CSF levels, individuals deemed to have high Tau levels maintained levels of M1 post-synaptic proteins (CADM3, NEO1, NPTX1, CHGB, PCSK1, NEGR, L1CAM, PTPRN, CACNA2D, PAM, VEGFA, NBL1 etc.) compared to pre-symptomatic individuals with lower Tau levels (48). This observation is analogous to differences we see between African Americans and Caucasians with AD. M1 members VGF and NPTX2, strongly correlate to antemortem cognitive measures (49-51) and VGF and NPTX2 has been nominated as biomarkers of neurodegeneration (N) as CSF levels enhance prediction of MCI to AD (51-53). Collectively, this would suggest that a specific sub-group of individuals with AD, including African Americans, have a higher burden of neurodegeneration (N) despite low CSF Tau levels. Future studies that analyze these CSF changes in neuronal proteins and other module members longitudinally will be important to better resolve these changes by race throughout disease progression.

Although a strength of our study was the large number of African Americans included there are several limitations that should be noted. First, we acknowledge that many of the protein changes we observe in the CSF across race could be due to ancestral or genetic differences (54,55). There was no genetics *a priori* performed on these study participants to confirm enrichment of African *vis a vis* European ancestry (56) as we stratified race solely by self-identification. Future studies, which include the integration of genetics and protein abundance to define protein quantitative trait loci (pQTL) will be necessary to resolve which proteins are under genetic control across race (57). Only a few studies to date have investigated proteomic difference by race in AD (58,59), which have predominately focused on brain tissues and not on the scale of this current study. However, a major initiative of the Accelerating Medicine Partnership (AMP)-AD partnership (60) is to increase the number of diverse tissues in the multi-omic analyses performed, which will complement data generated from these previous studies. Second, despite the well documented differences in the quality of education, higher rates of poverty, and greater exposure to adversity and discrimination that increase risk for dementia (4,5), these metrics were not captured on the participants in this study. Integrating CSF protein levels with vascular risk factors, and other environmental metrics may help better resolve some of the underlying racial differences in the CSF proteome. Finally, in this study we adjusted for co-factors such as age and sex to pinpoint changes that are most likely to be associated with race and AD. Sex and age have an impact on the abundance of CSF Tau and other protein levels (61). Therefore, future studies that assess the interactions between, age, sex and race will be informative. Nevertheless, this study reveals an impressive view of protein co-expression in AD CSF across race, which provides new insights into the pathways underlying cell-type changes and further evidence that race may mediate these in AD. These data also provide a resource for further investigations into how AD biomarkers vary across different racial backgrounds.

## Acknowledgements

This study was supported by the following National Institutes of Health funding mechanisms: U01AG061357 (A.I.L and N.T.S), R01AG070937-01 (J.J.L) and P30AG066511 (A.I.L) and the Foundations for the National Institute of Health AMP-AD 2.0 grant. We acknowledge Dr. Lenora Higginbotham and our colleagues at Emory for providing critical feedback.

## SUPPLEMENTAL FIGURES LEGENDS

**Supplemental Figure 1.**
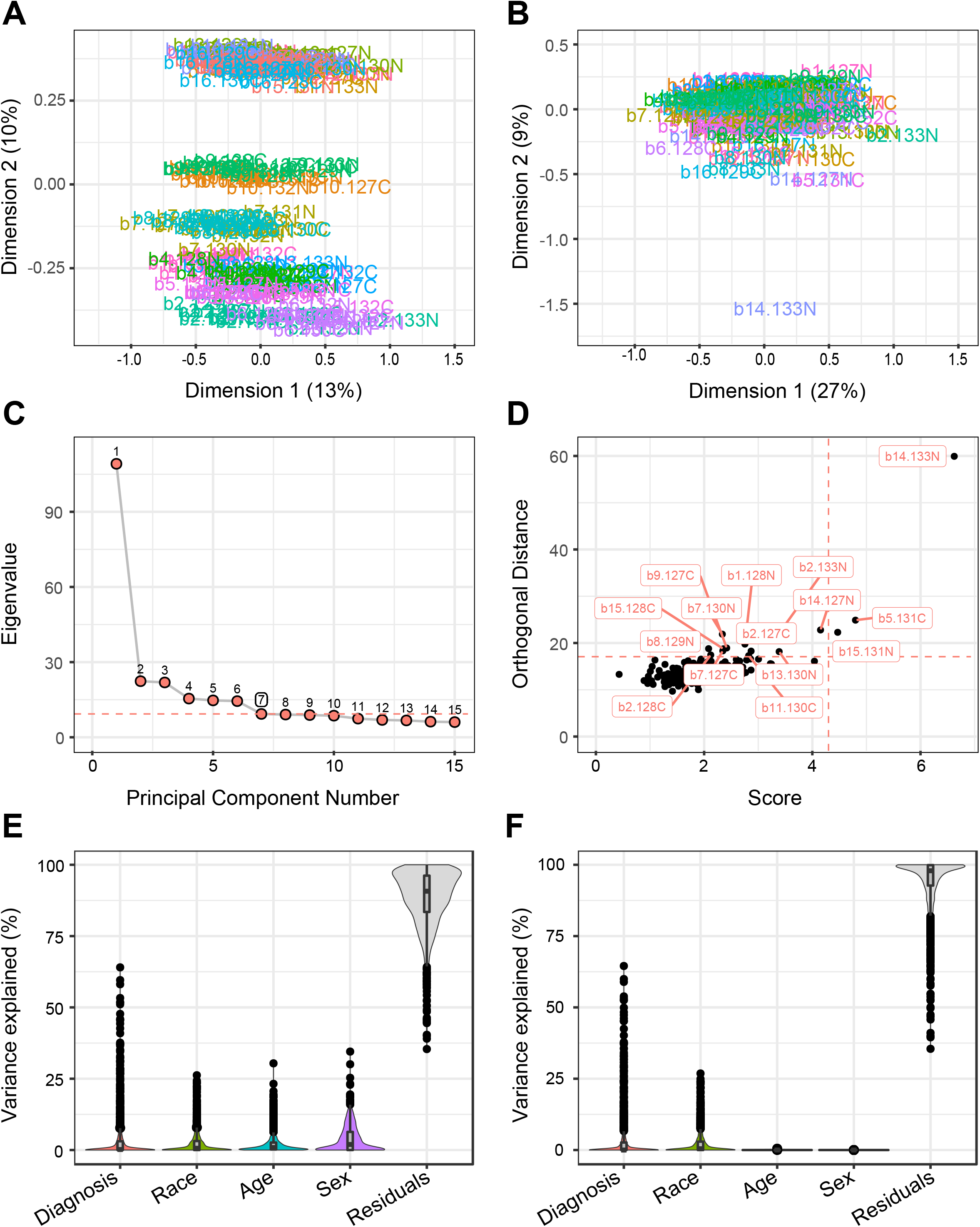
Batch correction, outlier removal and bootstrap regression. Multidimensional scaling (MDS) plots were used to illustrate batch contributions to variance before and after batch correction. In MDS plots, the distance a case is from one another is reflective of how similar or dissimilar a case is from the other. (**A**) Prior to batch correction, the samples clustered by batch (**B**) After batch correction, the samples no longer cluster by batch. (**C**) After batch correction, a principal component (PC)-based outlier removal method was utilized to detect outliers. By graphing the eigenvalue of each component against the PC number, the elbow or bend in the graph, which in this case was 7, was indicative of the ideal number of components to include within the parameters. (**D**) With a criterion for computing cutoff values set to 0.99, the cutoffs for the detection of outliers for the orthogonal distance and score were 16.79257 and 4.654674 respectively. This resulted in the detection of 15 outliers (b1.128N, b11.130C, b13.130N, b14.127N, b14.133N, b15.128C, b15.131N, b2.127C, b2.128C, b2.133N, b5.131C, b7.127C, b7.130N, b8.129N, b9.127C). B14.133N was such an extreme outlier because it was an empty channel. (**E**) After outlier removal, the matrix underwent bootstrap regression to remove variations in the dataset that were due to age and sex. Variance partition plots were employed to illustrate the percent contribution of diagnosis, race, age, and sex to the variance of each protein. (**F**) Following bootstrap regression, variations explained by age and sex were removed.

**Supplemental Figure 2.**
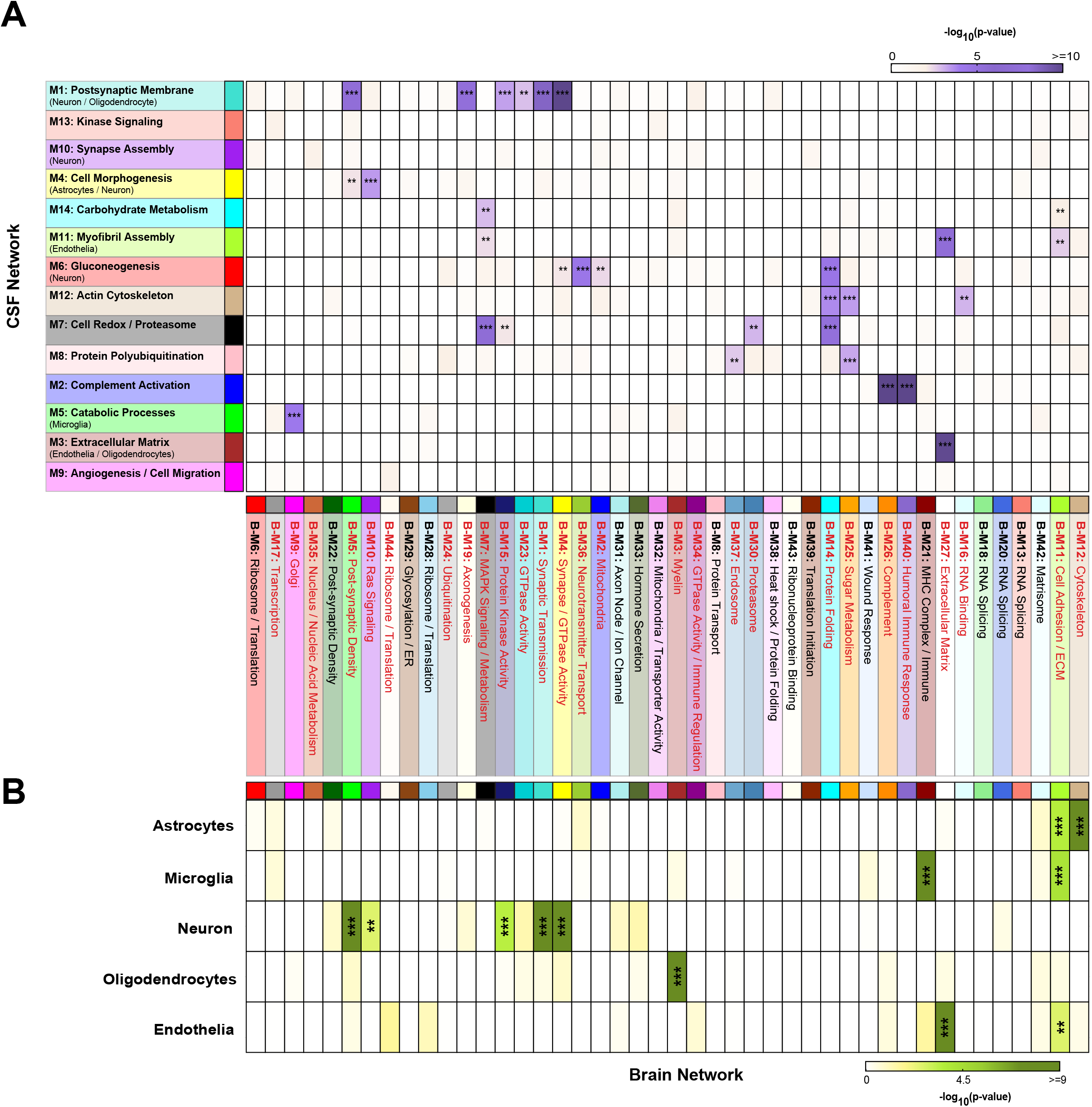
Protein overlap between modules in CSF network and modules in a human AD brain network. (**A**) Protein module enrichment across the CSF and brain was assessed by matching gene symbols of proteins in each module from the CSF network against gene symbols for protein in each module from a human AD consensus brain network using a one-tailed Fisher’s exact test The degree of enrichment increases from pink to light purple to dark purple with asterisks denoting the following statistical significance (**p≤0.01 and ***p≤.001). (**B**) Similar to CSF, cell-type enrichment was assessed by cross referencing brain module proteins against a list of proteins determined to be enriched in neurons, oligodendrocytes, astrocytes, and microglia using a one-tailed Fisher’s exact test. The degree of cell-type enrichment increases from yellow to green-yellow to dark green with asterisks denoting the following statistical significance (**<p≤0.01 and ***p≤.001).

**Supplemental Figure 3.**
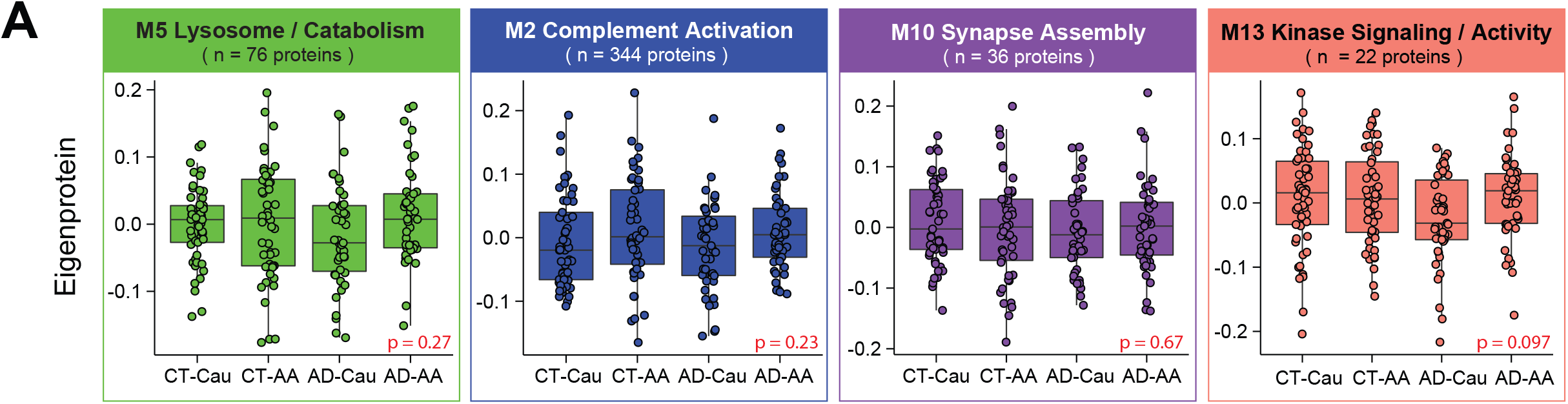
Additional CSF network protein modules. (**A**) Eigenprotein levels were distributed by race and diagnosis for remaining modules not shown in main Figure 4. This includes M5, M2, M10, and M13.

**Supplemental Figure 4.**
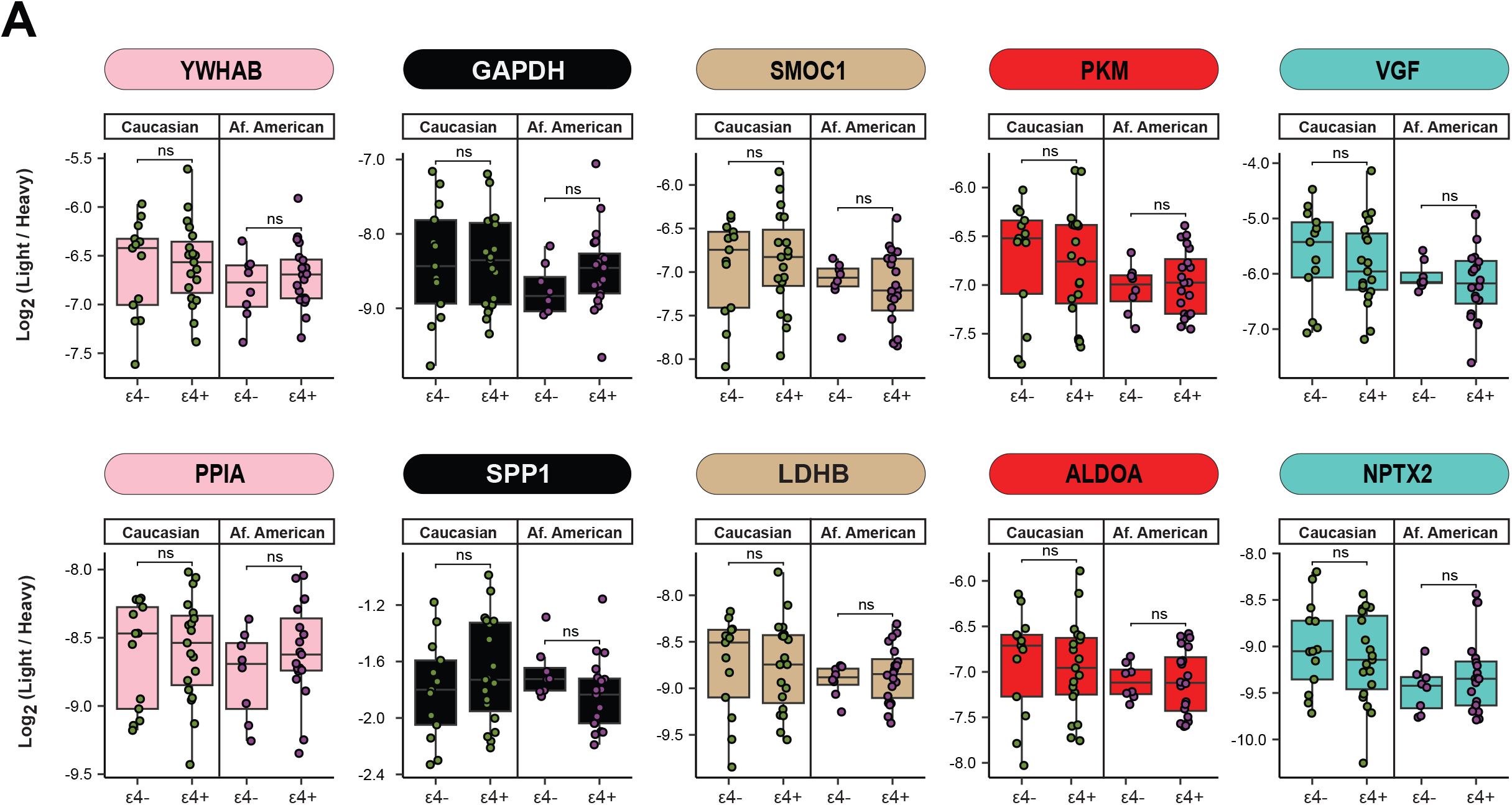
Stratification of SRM CSF protein measurements in AD by APOE genotype. (**A**) Within each race, protein abundances for were not affected by APOE ε4 genotype for YWHAB, GAPDH, SMOC1, PKM, VGF, PPIA, SPP1, LDHB, ALDOA, and NPTX2.

